# Bipartite functional fractionation within the default network supports disparate forms of internally oriented cognition

**DOI:** 10.1101/864603

**Authors:** Rocco Chiou, Gina F. Humphreys, Matthew A. Lambon Ralph

## Abstract

The ‘default network’ (DN) becomes active when the mind is steered internally towards self-generated thoughts but turns dormant when the mind is directed externally towards the outside world. While hypotheses have been proposed to characterise the association and dissociation between different component areas of the DN, it remains unclear how they coalesce into a unitary network and fractionate into different sub-networks. Here we identified two distinct subsystems within the DN – while both subsystems show common disinterest in externally-oriented visuospatial tasks, their functional profiles differ strikingly according to the preferred contents of thoughts, preferred modes of task requirement, and causative neural dynamics among network nodes. Specifically, one subsystem comprises key nodes of the frontotemporal semantic regions. This network shows moderate dislike to visuospatial tasks, shows proclivity for task-contexts with restraints on thoughts and responses, and prefers thoughts that are focused on other people. By contrast, the other subsystem comprises the cortical midline structure and angular gyri. This network shows strong aversion to visuospatial tasks, favours task-contexts allowing free self-generated thoughts without constraints, and prefers thoughts that are focused on self. Furthermore, causative connectivity reveals that task-contexts systematically alter the dynamics within and between subsystems, suggesting flexible adaption to situational demands. This ‘self/inward *vs.* others/outward’ separation within the broad DN resembles recent discoveries regarding a dyadic structure within the frontoparietal network that comprises regions controlling memories/thoughts *vs.* regions controlling sensory-motoric processes, and echoes burgeoning views that the brain is organised with a spectrum-like architecture along gradational changes of ‘inward *vs.* outward’ preferences.

**Significance:** Rather than construing the default network (DN) as ‘task-negative’ regions that passively react to off-task mind-wandering, researchers have begun to acknowledge the active role of the DN in supporting internally-directed cognition. Here we found a striking dichotomy within the DN in terms of the subsystems’ task-driven functional and connectivity profiles, extending beyond previous inferences using meta-analysis and resting-state fMRI. This dichotomy reflects a local manifestation of a macro-scale gradient representation spanning across the broad cerebral cortex. This cortical gradient increases its representational complexity, from primitive sensory and motoric processing, through lexical-semantic codes for language tasks, to abstract self-generated thoughts in task-free contexts. These findings enable a framework where the separate yet related literatures of semantic cognition and default-mode processes converge.

---

The discovery of the brain’s default network (DN) epitomises the serendipity of science. In the fledging period of human functional neuroimaging, researchers primarily employed externally-oriented tasks (e.g., visual search, speech recognition, or finger tapping) to demarcate and catalogue brain regions responsive to external stimuli. The serendipitous discovery of the DN is consequent to using passive viewing (rest) as a contrasting baseline – some early studies observed that a set of brain regions, including the angular gyri and midline structures (medial prefrontal and posterior cingulate cortices), reliably exhibit heightened activity during passive moments yet become deactivated during active tasks (e.g., 1). These task-negative resting-state activity was initially treated as an inadequacy of experimental design that failed to control for confounding variables (for discussion, see 2). The turning point in the field’s conceptualisation about the DN’s functionality came when Raichle *et al.* published their trailblazing work in 2001 – rather than treating the DN’s activity as a failure of designing a proper baseline, Raichle *et al*. postulated that the task-negative regions collectively contribute to human cognition when the mind is disengaged from external tasks and returns to the ‘default mode’ (rest) (3). After their seminal work, subsequent research found that the ebb and flow of neural activities in the DN regions tend to synchronise, forming a cohesively oscillating network (e.g., 4). It has been established that the DN amplifies its activity during various introspectively-directed tasks (e.g., reminiscing the past, imagining the future, reflecting on self, and empathising with others) (5-7). Such observations argue against defining the DN based on its absence during externally-directed tasks and instead suggest defining it using its active participation during introspection. While much research has been spawned since the early studies reviewed above, the complicated relationship between the DN and different types of introspective processes remains ill-understood. This background provides the milieu of the present investigation.

Although it is now apparent that the once prevailing ‘task-rest’ dichotomy is inaccurate and the ‘internal-external’ dichotomy more appropriately define the DN’s functionality, we still lack an encompassing framework to account for its omnipresent involvement in different types of introspective activities, as well as the nuanced subdivision within this network in relation to different tasks. To tackle these challenges, various attempts have been made (*i*) to parcellate the brain into functionally distinct networks and (*ii*) to uncover the cardinal organising principles with which brain regions join forces or split up. For instance, based on the brain’s task-free intrinsic connectivity and hierarchical clustering, it has been shown that the whole-brain’s inherent connectivity fractionates into (at least) seven primary networks (8), and the DN could be further partitioned into multiple sub-networks with them preferentially associated with different tasks (9). While such evidence demonstrates the heterogeneity of functional brain modules both at macro-scale (whole-brain) and meso-scale (within the DN), it remains unclear whether there is a cardinal organising dimension along which networks serving similar purposes unite while networks serving distinct purposes bifurcate. However, some recent studies offer clues that this organising dimension exists and, more importantly, operates as a general principle at multiple levels: First, Margulies *et al.* showed that the whole-brain’s complex connectivity pattern can be condensed into a topographical principle that explains a large proportion of cerebral layout, with regions serving outwardly-oriented activities (perception and action) on one end of the spectrum and regions serving introspective activities on the other end (10). A similar structure, albeit more rudimentary, is found in marmoset monkey’s brain, suggesting a shared evolutionary origin (11). Second, Vidaurre *et al.* investigated how resting-state activity unfolds in time and uncovered a systematic temporal structure that could be delineated as switching between two states – the propagation of neural activity tends to cycle within either the ‘outwardly-leaning’ sensorimotor network or the ‘inwardly-leaning’ default network, with sporadic transition between the two major networks (12). Third, Dixon *et al.* showed that the frontoparietal control network fractionates into two major subsystems with each involved in different types of executive control – one is more closely aligned with sensorimotor regions and more active when attending external stimuli whereas the other is more coupled with the DN and more active when attending to internal thoughts (13). In light of these recent discoveries, we asked a critical question – while the DN, as a whole, is more active for introspective processes, it remains unknown whether there exists a similar bipartite split within the DN, with one subsystem favouring more externally-oriented thoughts/contexts (e.g., empathising with other people) while the other preferring more internally-oriented thoughts/contexts (e.g., contemplating about self).

The research literature of semantic cognition offers important clues as to how the DN might fractionate into subsystems. Semantic cognition refers to the high-order human capacity to comprehend the meaning of inputs (e.g., to understand text/speech, to recognise a corkscrew and its function) and produce behavioural outputs according to meaning (e.g., to correctly use a corkscrew with twisting action) (14). It is a critical cognitive faculty that interfaces internal representations of meaning (all our pan-modality knowledge about the world) with external modalities (all the incoming auditory, visual, tactile signals that need to be mapped onto generalisable, modality-invariant concepts). Decades of research have shown that the two major aspects of semantic cognition – the ability to represent contents of semantic knowledge and the ability to manoeuvre semantic contents in appropriate ways – rely on different nodes of the frontotemporal semantic network (SN). Semantic knowledge *per se* is encoded primarily by the anterior temporal lobe (ATL) and various auxiliary regions (15), whereas the ability to flexibly select and recombine semantic knowledge relies jointly on the inferior frontal gyrus (IFG) and posterior middle-temporal gyrus (pMTG) (for reviews, see 14, 16, 17). Intriguingly, previous research on default-mode processes often incorporates these SN regions – the ATL, IFG, and pMTG – as the *extended* prongs of the DN, as compared to the three *core* regions of the DN – the medial prefrontal cortex (mPFC), posterior cingulate cortex (PCC), and angular gyrus (AG) (e.g., 9, 18, 19). Incorporating the SN into the DN is due to their commonality: just as the core DN nodes, the key SN nodes are situated within the cortical realm that is deactivated by externally-directed non-semantic tasks (particularly the ATL, see 20). Moreover, the SN regions have been found to link robustly with a broad swathe of the DN during both task- and resting-state (18, 21, 22). There are, however, differences – it has been shown that while the ATL and AG are both deactivated by externally-directed visuospatial tasks, they show opposite patterns to semantic processes, with ATL activity elevating while AG activity lowering for semantic tasks (23). Taken together, these commonalities and differences motivate our focus on exploring the relationship between the DN and SN, with the aim to understand the division within the broad DN territory and the neurocognitive dimension behind the fractionation.

In the present study, we report findings of two fMRI experiments whereby we systemically manipulated two critical neurocognitive dimensions that regulate the DN’s activity: First, participants were either constrained by specified correspondences among external stimuli, semantic meaning, and reaction within a short time-frame (Experiment 1) or were allowed to let internal theme-guided thoughts roam with little time pressure (Experiment 2); the former context demands attention to the external entities (visual stimuli and effector reaction), whereas the latter emphasises attention to internal thoughts. Second, cognitive operations were directed towards visuospatial processing of external meaningless stimuli (mental rotation or visual search), internal thoughts about self (reflecting on self-traits, or autobiographical memory), internal thoughts about others (reflecting on others-traits, or mentalisation), as well as task-free mind-wandering (rest). To pre-empt the main findings, with these systematic manipulations, we found a clear bipartite split within the DN. While the broad DN, as a whole, strongly prefers introspective activity to perception and action, it fractionates functionally into two bodies – one subsystem favours self-referential processing and is tempered by externally-directed task-constraints, whereas the other subsystem favours others-referential processing and is reinforced by external task-constraints. Moreover, neural dynamics within and between the two subsystems alter according to whether the task-contexts accentuates external stimuli or internal thoughts, providing further support to the bipartite organisation. These findings are discussed with reference with recent theories of the macro-scale transition from unimodal/sensorimotor to transmodal/abstract processing that spans across the entire human cerebrum.

## Results

Experiment 1 created a restrictive and relatively outwardly-directed context wherein participants read text about personality traits, assessed its pertinence with regard to either self or another individual, and made a response within a short timeframe using specified rules about the mapping between semantic meaning and manual responses (see *Methods* and *SI*). By contrast, Experiment 2 created a less restrictive and inwardly-directed context wherein participants were asked to recall autobiographical memory (AM) following a cue-photograph’s theme, imagine how the people in the photograph might be thinking or feeling (theory of mind/ToM). No response restriction and little time pressure were imposed while participants engaged in AM and ToM, allowing the mind to meander freely around thoughts/memories related to the tasks. Analyses were performed at multiple scales: We first interrogated the entire brain to examine whether differential brain structures exhibit preferences for distinct types of introspective activities. We then zoomed in on the regions of interest (ROIs) within the DN and SN, comparing the differential functional profiles of the two systems, using independently selected ROIs based on meta-analysis outcomes (details in *Method*s). In addition to the direct contrast between DN and SN nodes, we focused specifically on the functional sub-division at a provincial-scale, inspecting the responses of adjacent areas within the mPFC and within the inferior parietal lobule (IPL). This local-scale inspection was motivated by recent mounting evidence that (*i*) the broad mPFC zone harbours two sub-sections, with its dorsomedial part (the dmPFC) being more connected with the SN and its ventromedial part (the vmPFC) being more connected with the DN (18), and that (*ii*) a similar bisection has also been observed in the IPL, with its anterior section (the temporoparietal junction/TPJ) being more connected with the SN and posterior section (the AG) more connected with other DN nodes (24, 25).

### Mind-wandering (rest) activity reveals a bipartite split between the DN & SN

The traditional definition of the default mode system is a set of brain regions whose activity drops when the mind is engaged by externally-oriented tasks yet rises when the mind returns to the resting/default state. While the DN and SN both fall within the realm of this purported ‘task-negative’ zone, it remains unclear whether they respond differentially to mind-wandering (rest) *vs.* externally-oriented visuospatial tasks. To this end, we included, in both experiments, visuospatial control tasks that are design to elicit deactivation of the classic ‘task-negative’ zones – in Experiment 1, participants mentally rotated scrambled stimuli and compared their visual configurations, while in Experiment 2 they searched for a tiny triangle hidden in mosaic patterns (see *Methods*). We began by a whole-brain interrogation to identify regions that are significantly more active during rest, compared to the visuospatial conditions, stringently thresholded using FWE-correction (*p* < 0.05) for whole-brain voxel intensity. This search corroborates the classic definition of the ‘task-negative’ neuroanatomy: As illustrated in Fig. 1*A*, an extensive swathe of the DN (peaking at the bilateral mPFC, PCC, and AG), as well as the SN (peaking at the left ATL, IFG, and pMTG), robustly amplify their activity during rest, compared to the visuospatial tasks, consistently observed in both experiments (also see *SI* for the ‘task-positive’ regions that heightened activity for visuospatial tasks, relative to rest). Importantly, brain regions belong to the DN exhibit strikingly greater responses to the mind-wandering state (as indicated by greater magnitude of deactivation for the visuospatial tasks), relative to the SN regions. This is observed both at the regional and multi-node network levels: At a local level (within the mPFC), mind-wandering triggers significantly greater activity of the vmPFC (a sub-region more closely linked with other DN regions) compared to the dmPFC (a sub-region more closely linked with the SN), resulting in reliable effects seen in both experiments (*F*_(1,23)_ = 13.16, *p* = 0.001, *η*_p_^2^ = 0.36; see Fig. 1*B*). At a network level (the broad ‘task-negative’ zones), the three ROIs known to form the core DN (the vmPFC, PCC, and AG; see 9) show significantly greater activity for mind-wandering, relative to the three ROIs known to form the core SN (the ATL, IFG, and pMTG; see 14), also reliably found in both experiments (*F*_(1,23)_ = 56.20, *p* < 0.001, *η*_p_^2^ = 0.71; see Fig. 1*C*). Taken together, despite the fact that the DN and SN both show a tendency to turn active during resting-state, the DN regions, as a whole, respond much more intensively to mind-wandering, dwarfing the SN regions that respond more moderately. Moreover, our results indicate that while the wide mPFC is sometimes conflated as a unitary, monolithic region, it is actually heterogeneous, with its ventral section leaning towards the DN and responding vigorously to mind-wandering and its dorsal section leaning towards the SN and responding moderately.

**Fig. 1.**
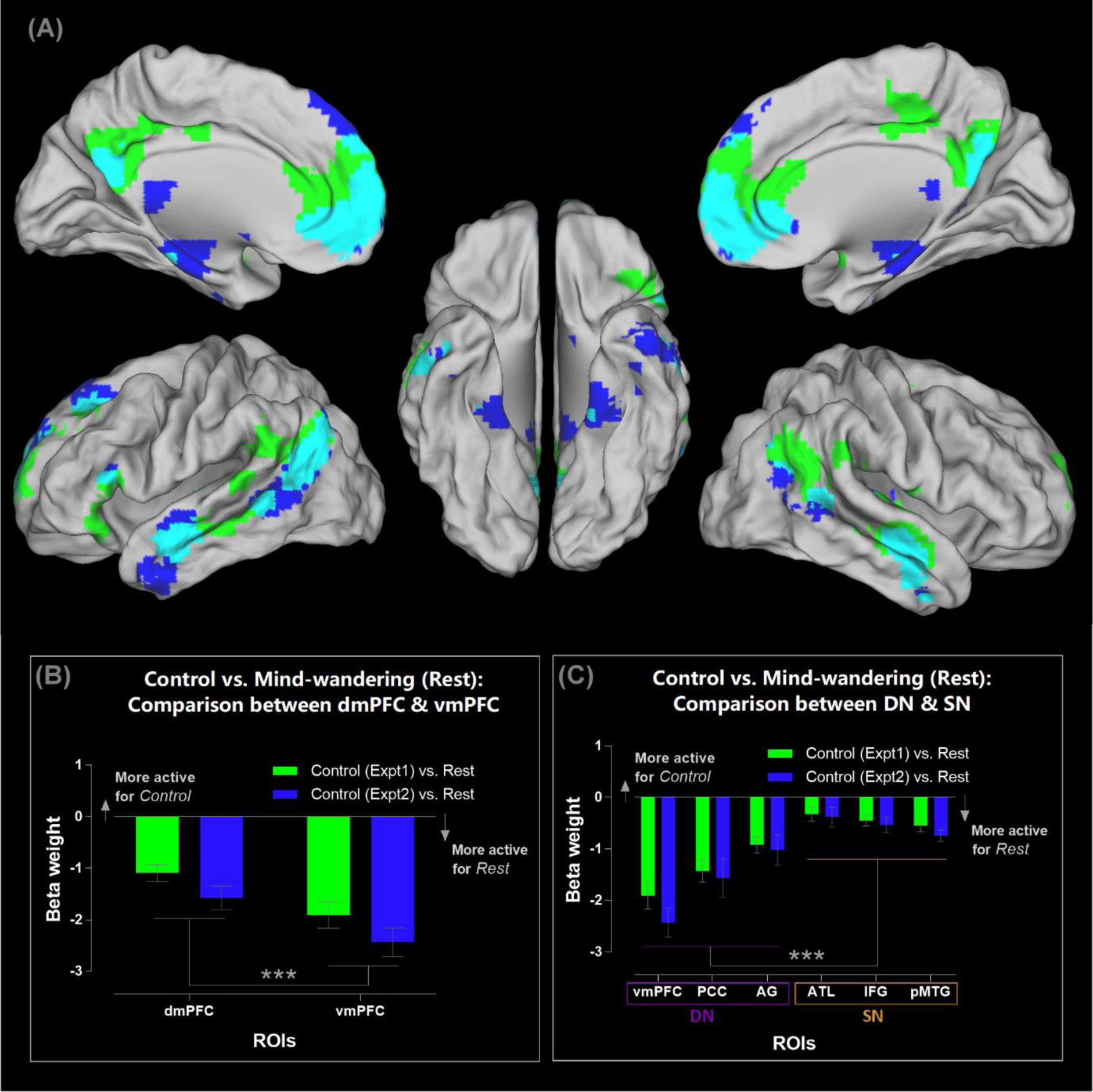
**(A)** Brain regions showing significantly greater activity for mind-wandering (rest), compared to the visuospatial control task, thresholded at *p* < 0.05 (FWE-corrected for whole-brain voxel intensity). Significant clusters in Experiment 1 (green), Experiment 2 (blue), and their conjunctions (cyan). **(B)** ROI-analysis of the dmPFC *vs*. vmPFC. Note that negative beta weight means *more active* for mind-wandering (rest), compared to the visuospatial control tasks. **(C)** ROI-analysis of the three core DN areas (the vmPFC, PCC, and AG), as well as the three core SN areas (the ATL, IFG, and pMTG). See *SI* for the locations of ROIs that are rendered on a template brain. *** *p* < 0.001

### Different types of self-referential mental activities reveal a bipartite split between the DN & SN

Mental activities related to ‘self’ are intuitively ‘inwardly-leaning’ and have been known to strongly associated with the DN (for review, see 26, 27). In particular, much emphasis has been laid on the mPFC – various types of self-related mental activities robustly engage this area, and pathology of the mPFC leads to difficulty in self-regulation (for review, see 28). Previous neuroimaging investigation into the neural basis of self-related processes have adopted two primary types of experimental approaches: One pervasive approach is asking participants to access self-concept via evaluating adjectives about personality traits with reference to self, which entails participants processing semantic information, bearing the task rules in mind, and reacting on a trial-by-trial basis within a time limit (e.g., 7, 29). The other popular approach is asking participants to recollect autobiographical memories related to a provided topic while participants have sufficient time (typically an interval longer than 10s) to engage in mnemonic retrieval (e.g., 5). The former approach is more ‘semantic’ in nature and entails an outwardly-focused context (reading text, pushing buttons), whereas the latter is more ‘episodic’ in nature and entails an inwardly-focused context. Our experiments provided us a unique window into how the DN and SN might respond differently to self-related activities under these two types of situations. We began unpicking this difference by examining the whole-brain activity pattern, comparing the distribution of activities induced by the self-traits assessment task (‘semantic-self’) and the autobiographical recollection task (‘episodic-self’). As illustrated in Figure 2*A*, both tasks significantly increase activities of various DN and SN regions compared to rest, replicating previous data that goal-directed introspective activities heighten the DN compared to aimless mind-wandering (5). However, closer inspection on the pattern reveals that while expansive swathes of the SN regions are recruited by both tasks, the three core DN nodes (the vmPFC, PCC, and AG) are exclusively recruited only when one processes ‘episodic-self’ during the autobiographical task. Such differential engagement of the DN and SN in the two types of self-related processing becomes clearly manifested when we examine the ROIs (note that all of the contrast baseline here is rest/mind-wandering, rather than external-visuospatial tasks, hence allowing us to assess whether goal-directed introspective activity further enhances neural activity, without the contamination from the suppressive effects of external tasks): As Figure 2*B* shows, within the mPFC, its ‘SN-leaning’ dorsal section is more active for self-concept, whereas its ‘DN-leaning’ ventral section is more active for autobiographical memory, resulting in a significant interaction (*F*_(1,23)_ = 29.37, *p* < 0.001, *η*_p_^2^ = 0.56). This dissociation is not only found at the local scale within the mPFC, but also seen at a more global scale: As Figure 2*C* illustrates, while nearly all of the ROIs exhibits preference for autobiographical memory over self-concept, this preference is evidently much attenuated in the three ROIs belong to the SN, particularly in the inferior frontal gyrus (*F*_(1,23)_ = 19.72, *p* < 0.001, *η* _p_^2^ = 0.46). To dissect this interaction further, we separately scrutinised the neural response to each task and found that retrieval of autobiographical memory engages regions of the DN and SN to an equivalent extent (*p* > 0.32, Fig. 2*D* right). By stark contrast, accessing self-concept primarily recruits the SN yet minimally engages the DN, leading to a significant difference (*p* < 0.001, Fig. 2*D* left). Finally, we plotted the magnitude of preference for autobiographical memory over self-concept (indexed as their *β*-weight difference) for each ROI to manifest their relationship: As Fig. 2*E* shows, while all of the ROIs are more active for autobiographical recollection, there is a clear split, with all of the DN nodes exhibiting strong preference *above* the average and median level (driven primarily by their disengagement during the self-concept task) and all of the SN nodes showing much mitigated preference falling *below* the average and median (indicating the SN’s prevalent participation in both tasks). Taken together, these results clearly indicate a bipartite split in the neurocognitive bases for different types of self-related processing – accessing self-related semantic concepts relies on the SN while barely involves the DN (see *SI* for further discussion regarding how the choice of contrast baseline affects whether the mPFC is revealed in the ‘self-traits’ type of tasks), whereas accessing episodic memories of self-experiences is a comparatively more engaging process that recruits both the DN and SN.

**Fig. 2.**
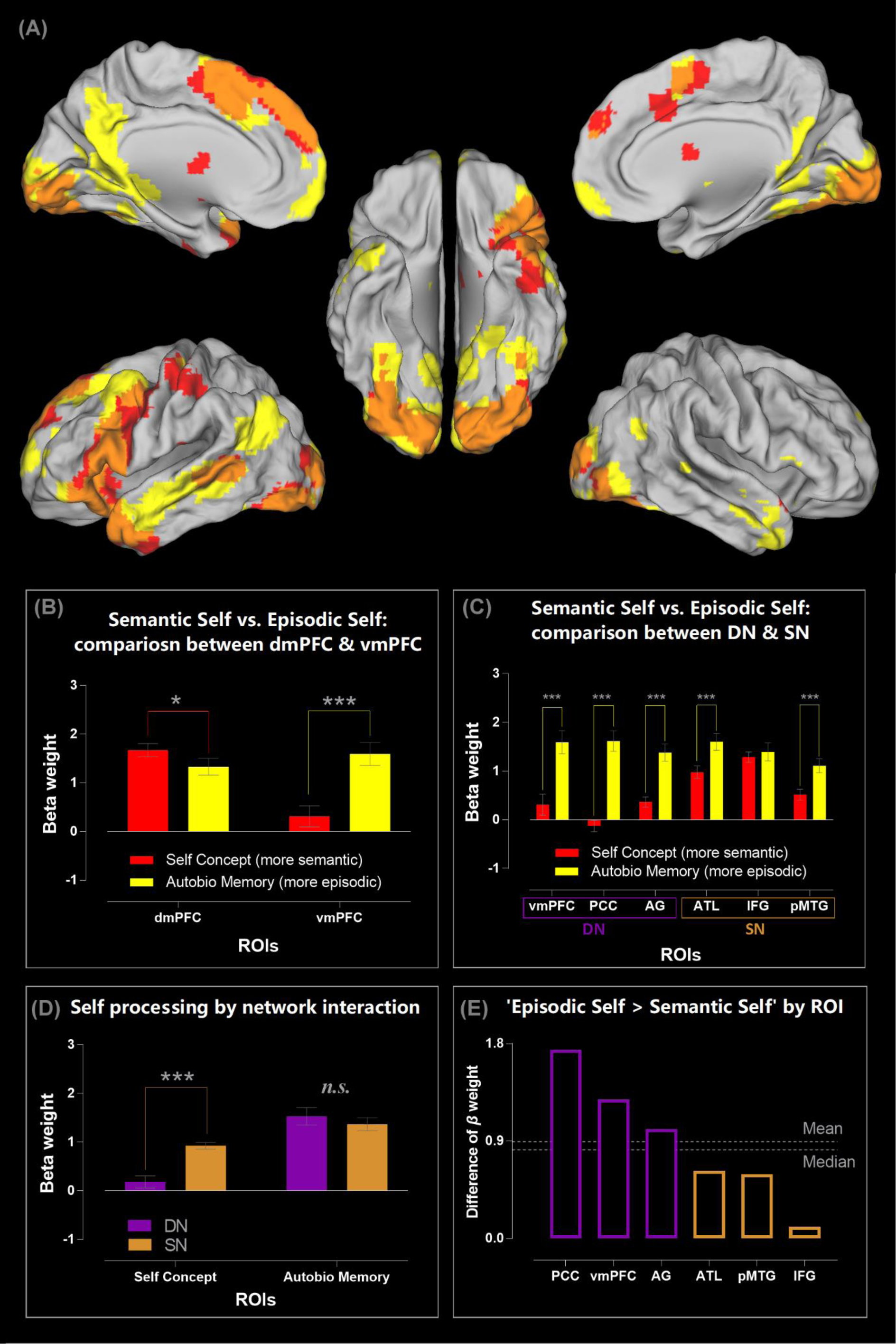
**(A)** Brain regions showing significantly greater activity for the two types of self-related tasks: the self-concept evaluation task (red), for the autobiographical memory task (yellow), and their conjunctions (orange). All contrasts were against the baseline of mind-wandering (rest), thresholded at *p* < 0.05 (FWE-corrected for whole-brain voxel intensity). **(B)** ROI-analysis reveals an interaction between brain regions (dmPFC *vs*. vmPFC) and the types of self-related tasks (self-concept *vs*. autobiographical memory). **(C)** ROI-analysis of the three core DN areas (the vmPFC, PCC, and AG), as well as the three core SN areas (the ATL, IFG, and pMTG). **(D)** The significant interaction between the two networks and the two types of self-processing. **(E)** The six ROIs ranked by their preference for autobiographical memory over self-concept, with all of the three DN regions showing a noticeably greater preference than all of the three SN regions. *** *p* < 0.001, * *p* < 0.05

### Self- *vs.* other-referential mental activities reveal a bipartite split between the DN & SN

Decades of neuroimaging research has accumulated a myriad of evidence for a common neural basis for understanding the mental states (including decisions, feelings, beliefs) of both self and others (30-32). This shared neurocognitive system for both self- and other-referential processing comprises the cortical midline-structures (i.e., the dmPFC, vmPFC, and PCC) and some lateral regions (including the ATL, IFG, pMTG, and IPL). However, it remains unclear whether the constituent regions in this system split into subsystems depending on whether they prefer inwardly- (e.g., reflecting on self) *vs*. outwardly-directed (e.g., mentalising about others) socio-cognitive processes. Experiment 2 provided conducive conditions to study whether there is a schism due to preferred processes – during the AM situation, participants recollected their own thoughts and feelings associated with specific autobiographical events, whereas during the ToM situation, they made inferences about the mental states of other people. While the two tasks differed on the focus of social target (inward/self *vs*. outward/others), they were yoked and matched by our experimental design: Identical, counterbalanced stimuli were used in both conditions to rule out stimuli-related effects, and no external response was required during the AM and ToM periods (see *Methods*) to encourage participants to concentrate on introspective experiences during the lengthy interval. Results of the whole-brain search, based on the direct contrast between the AM and ToM contexts, reveal a clear bipartite split that corroborates our speculation: As shown in Fig. 3*A*, all of the three core DN regions (the vmPFC, PCC, and AG) were preferentially more engaged by AM compared to ToM. By contrast, extensive stretches of the SN regions (including the three core SN nodes – the bilateral IFG, ATL, and pMTG, as well as other semantic-related regions in the inferior temporal-parietal cortices) were more engaged by ToM than AM. Analysis of ROIs further highlights the granularity of such bipartite structure: As shown in Figure 3*B*, an evident split is found within the mPFC, with its ‘SN-leaning’ dorsal section preferring other-referential/ToM processes and its ‘DN-leaning’ ventral section preferring self-referential/AM processes (*F*_(1,23)_ = 15.87, *p* = 0.001, *η*_p_^2^ = 0.41). The split is also observed at the network level (Fig. 3*C*), with the core DN midline-structures (i.e., the vmPFC and PCC) exhibiting a strong preference for self-over other-referential processes and all of the three SN nodes (the IFG, ATL, pMTG) exhibiting a marked preference for other over self (*F*_(1,23)_ = 42.43, *p* < 0.001, *η*_p_^2^ = 0.65). Next, we focused on the IPL – this analysis is motivated by two threads of evidence: (*i*) the anterior IPL sector (i.e., the TPJ) is more closely linked with the ‘SN-leaning’ dmPFC whereas the posterior sector (the AG) is more linked with the ‘DN-leaning’ vmPFC (18); (*ii*) the bilateral TPJ has been shown to be involved in simulating the mental states of other people, with the right TPJ tending to show more pronounced responses than its left counterpart (e.g., 6, 33). We selectively examined the response profiles of the bilateral AG and TPJ, and the results are shown in Fig. 3*D*: Although the AG and TPJ are adjacent to each other, they prefer different types of introspective activities (*F*_(1,23)_ = 93.34, *p* < 0.001, *η*_p_^2^ = 0.80): The bilateral AG show an overall preference for the self-referential AM condition over the other-referential ToM condition (*p* = 0.007, although the preference is more exaggerated in the right AG). By contrast, the TPJ show an overall preference for ToM over AM, reliably seen in both hemispheres (*p* < 0.001). Finally, we focused on a set of regions displaying a preference for other-referential processing – the dmPFC and the bilateral TPJ. We specifically examined whether their response profiles alter with different types of other-referential processing: assessing the personality traits of another person (Experiment 1) *vs*. simulating what someone might be thinking or feeling (Experiment 2). A significant interaction statistically supports their distinct characteristics (*F*_(1,23)_ = 7.75, *p* = 0.001, *η*_p_^2^ = 0.25): While the response amplitude of the dmPFC does not differ between the personality knowledge and ToM contexts (*p* = 0.27; also see *SI* for supplemental results concerning the equivalent dmPFC involvement for the personality assessment of self *vs.* other), the left TPJ (*p* = 0.002) and right TPJ (*p* < 0.001) both reveal a robustly greater response to ToM compared to personality knowledge. Taken together, these analyses provide convergent evidence to our earlier findings – there is an ‘inward *vs*. outward’ principle that governs the neurocognitive architecture of the broad default-mode system, with the core DN regions preferring tasks encouraging introspective activity and a focus on self and the SN regions preferring tasks necessitating externally-directed activity (attending to outside stimuli and response) and a focus on others. Furthermore, our data corroborate a division of labour that has been documented previously (28, 30, 34) – within the subsystem that prefers other-referential processing, the dmPFC is recruited whenever a task involves ascribing socio-emotive features to an animate being, be it self or other, whereas the TPJ is exclusively recruited for ‘online’ stimulation of mental states during the ToM-type of tasks.

**Fig. 3.**
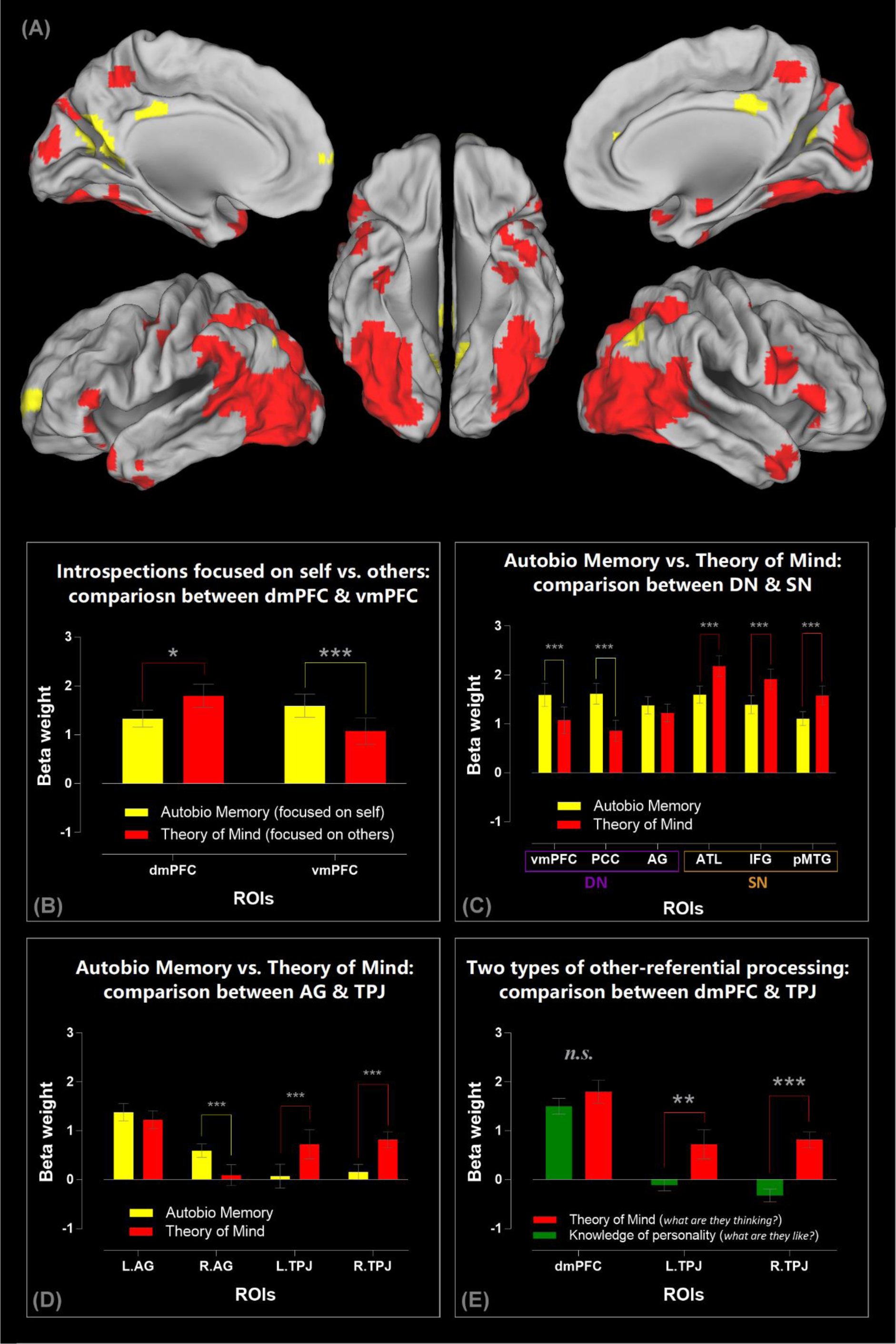
**(A)** A direct comparison between the two introspective tasks reveals the regions showing significantly greater activity for the autobiographical memory task (self-referential: yellow) and for the theory of mind task (other-referential: red). Statistics are thresholded at thresholded at *p* < 0.05 (FWE-corrected for whole-brain voxel intensity). **(B)** ROI-analysis reveals an interaction between brain regions (dmPFC *vs*. vmPFC) and introspective tasks (autobiographical memory *vs.* theory of mind). **(C)** ROI-analysis of the three core DN areas (the vmPFC, PCC, and AG), as well as the three core SN areas (the ATL, IFG, and pMTG). **(D)** The significant interaction between the two inferior-parietal subregions (the AG *vs.* TPJ) and the two types of introspective processing. **(E)** The significant interaction between the regions preferring other-referential tasks (the dmPFC, left TPJ, and right TPJ) and the two types of other-referential processes. *** *p* < 0.001, ** *p* < 0.01, * *p* < 0.05

### Externally- and internally-oriented task-focuses modulate the neural dynamics of the DN & SN

We capitalised on dynamic causal modelling (DCM; 35, 36) to investigate whether and how the causative/directional neural dynamics within and between the DN and SN alter as a result of switching between externally- and internally-directed task-focus. Decades of validations based on empirical fMRI data have substantiated the efficacy of DCM in making inferences about the causative connectivity between brain regions, outperforming other analytical approaches of causative connectivity (for review, see 37). Particularly, DCM enables researchers to apply a Bayesian statistical procedure to compare and select, amongst multiple candidate models, the ‘winner’ model that offers the most probable explanation regarding the mechanistic neural implementation that generates the observed fMRI data (38, 39). This statistical procedure estimates the ‘protected exceedance probability’ (39), indicating whether a model’s explanatory power surpasses any other candidate (as compared to the remaining models) above and beyond the likelihood of all models being equiprobable (as compared to chance). Here we report a series of DCM analyses, investigating whether the directionality of communication between/within the DN and SN changes depending on whether the task encourages mental activities that are more externally-oriented (Experiment 1: the personality-trait tasks that confine thoughts, require explicit motoric response, and stipulate stimulus-response mappings) *vs*. internally-oriented (Experiment 2: the AM and ToM tasks that allow unrestrained thoughts and do not entail any overt response). As discussed below (also see *Methods*), for different models we assumed that incoming signals enter the network through different brain areas (which belongs to either the DN or SN), which represents the ‘inception’ event that triggers subsequent neural dynamics. We then employed Bayesian statistical methods to identify the best model that underlies the fMRI data.

The first (Experiment 1; Fig. 4*A*) and second (Experiment 2; Fig. 4*B*) DCM analyses were focused on the mPFC. Our earlier analysis reveals a local-scale bipartite split, with the dmPFC favouring externally-directed tasks and the vmPFC favouring internally-directed tasks. Based on this, we examined whether task-context impact on the ‘starting-point’ from which the flow of neural processing cascades downstream. This was implemented by assuming, for different models, a different ‘entry’ location through which the triggering input enters the two-node network (dmPFC *vs*. vmPFC). For each DCM, we constructed three models, hypothesising that the trigger enters the system through the dmPFC, vmPFC, or both nodes (see *Methods* and Fig. 4*A* and 4*B*). Exactly identical coordinates of the dmPFC and vmPFC were applied to localise the network nodes for the analyses of Experiment 1 and 2. This permits a rigorous test on whether the very same group of brain regions adjust their interplay under externally- *vs*. internally-veered contexts, through the contrast of the two DCM outcomes. Using well-established Bayesian procedures (39, 40) to select the model that provides the best mechanistic explanation that underlies the observed data, we found that task-focus drastically changes the directionality of neural dynamics. As clearly indicated by the magnitude of protected exceedance probabilities, when the task focussed on externally-directed mental activities (Experiment 1; Fig. 4*A*), the dmPFC-input model overwhelmingly outperformed other models and was the *only* model that substantially exceeded the chance level. By contrast, when the task encouraged internally-directed processes (Experiment 2; Fig. 4*B*), the vmPFC-input model became the winner that strikingly outperformed other models and was the *only* model that performed above chance. This demonstrates the reversal effect of task-contexts – when the task was directed externally, the SN-leaning dmPFC initiated the onset of neural dynamics; by contrast, when the task was directed internally, the DN-leaning vmPFC became the trigger.

**Fig. 4.**
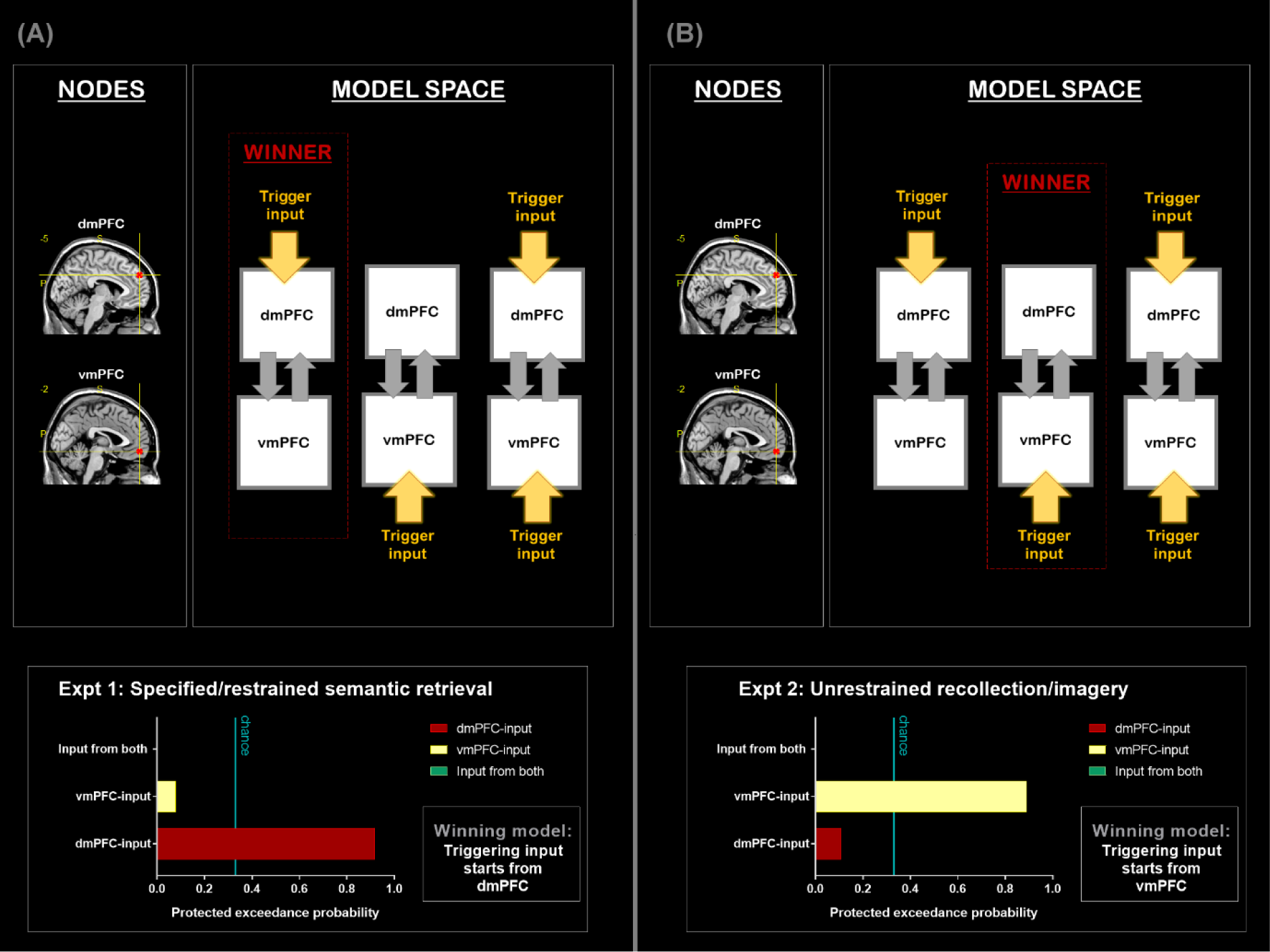
**(A)** DCM-1: All analyses are based on the data of Experiment 1; **(B)** DCM-2: All analyses are based on the data of Experiment 2. Note that the nodes of the two DCM have the same coordinates, suggesting a drastic change of network dynamics as a result of experimental contexts.

Next, in the third (Experiment 1; Fig. 5*A*) and fourth (Experiment 2; Fig. 5*B*) DCM, we focused on the trilateral communication among the IFG, the ATL, and the vmPFC. These rostral-frontotemporal nodes were included in the model owing to their respective roles in controlling semantic retrieval (the IFG: 41, 42), representing semantic concepts (the ATL: 14), and underpinning internally-focused thoughts (the vmPFC: 5). We constructed three models for each DCM, assumed a different starting-point for each model (IFG-input, ATL-input, or vmPFC-input), and set both DCM to be based on identical nodes. Results of Bayesian model selection are shown in Fig 5*A*: During the externally-focused Experiment 1, the IFG-input model exceedingly outperformed other models and was the *only* model that surpassed chance levels. However, during the internally-focused Experiment 2 (Fig. 5*B*), the network changed the dynamics between nodes – the vmPFC-model became the one that won over other models by a massive margin, offering the best account for the underlying neural interactions. This yields a consistent pattern across analyses – under an externally-biased context, neural dynamics stemmed from an SN site (DCM-1, DCM-3), whereas under an internally-biased situation, neural dynamics commenced from a DN site (DCM-2, DCM-4).

**Fig. 5.**
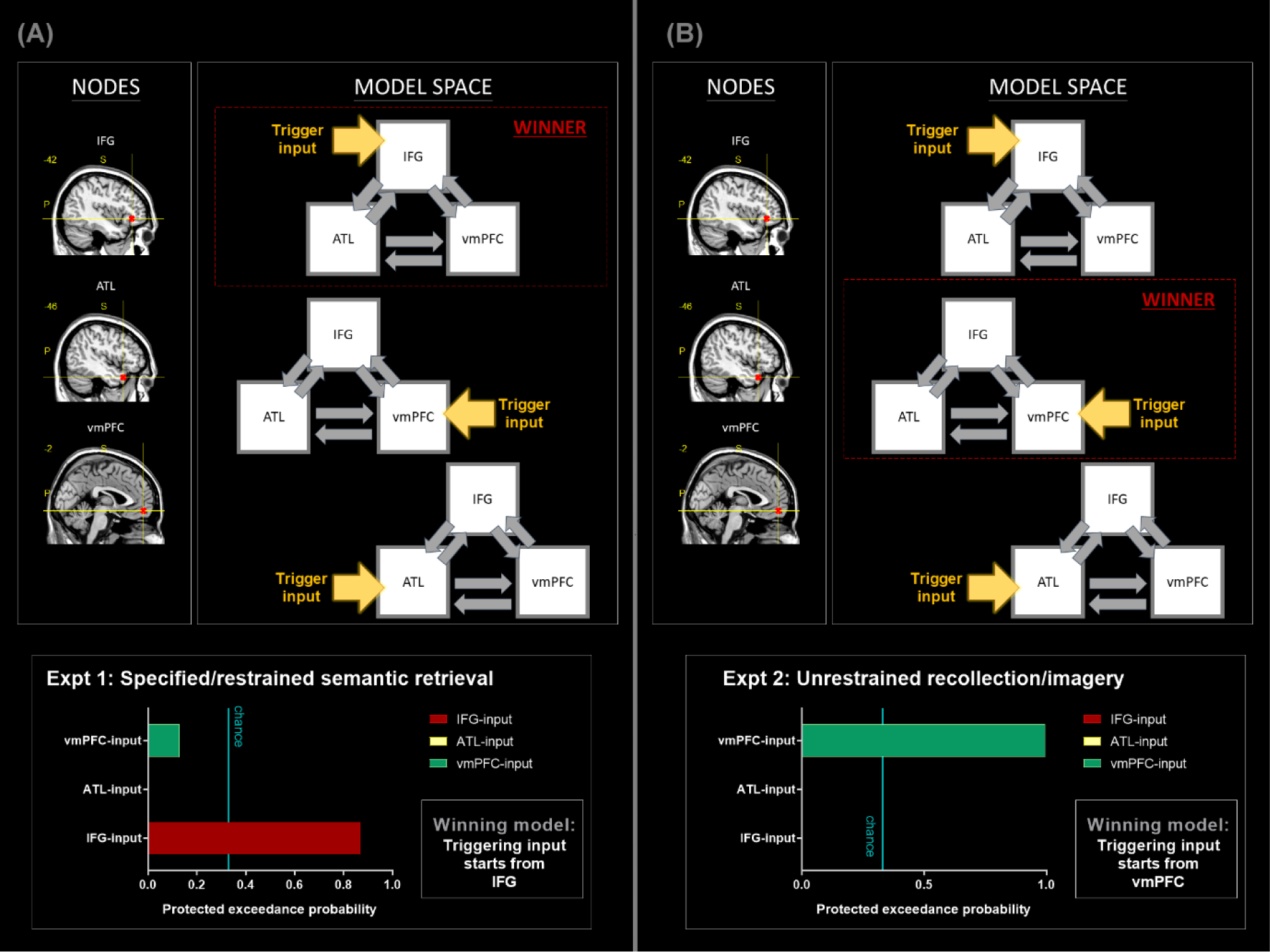
**(A)** DCM-3: All analyses are based on the data of Experiment 1; **(B)** DCM-4: All analyses are based on the data of Experiment 2. Note that the nodes of the two DCM have the same coordinates, suggesting a drastic change of network dynamics as a result of experimental contexts.

Finally, in the fifth (Experiment 1; Fig. 6*A*) and sixth (Experiment 2; Fig. 6*B*) DCM, we specifically focused on the SN for Experiment 1, and on the DN for Experiment 2, motivated by the robust contextual modulations that we observed in earlier analyses. These analyses aimed to identify, amongst all constituent sites within the SN/DC, the most reliable region that launches neural dynamics under an external-/internal-biased context. Thus, in DCM-5, we included all the key nodes closely associated with the SN – the IFG, the ATL, the pMTG, and the dmPFC. We constructed three models, hypothesising the triggering signal might start from the IFG, the pMTG, or the dmPFC. Results of Bayesian model selection showed that, as illustrated in Fig. 6*A*, the IFG-input model gained the highest protected exceedance probability and was the *only* model above the chance threshold, trumping other models. By contrast, in DCM-6, we included all the core nodes of the DN – the vmPFC, the PCC, the AG. Three models were built, with the triggering signal entering via the vmPFC, the PCC, or the AG. As shown in Fig. 6*B*, results of model comparison indicated that the vmPFC-input model gained most Bayes posterior likelihood, greatly outperforming other hypotheses. Taken together, this series of DCM analyses clearly demonstrated the fluidity of context-dependent neural dynamics – akin to a tug-of-war occurring at the neural level, an externally-focused task propels information to flow from an SN origin to DN (with the IFG being the most reliable triggering point), whereas an internally-focused task drives signal to travel from a DN origin to the SN (with the vmPFC being the most reliable triggering unit).

**Fig. 6.**
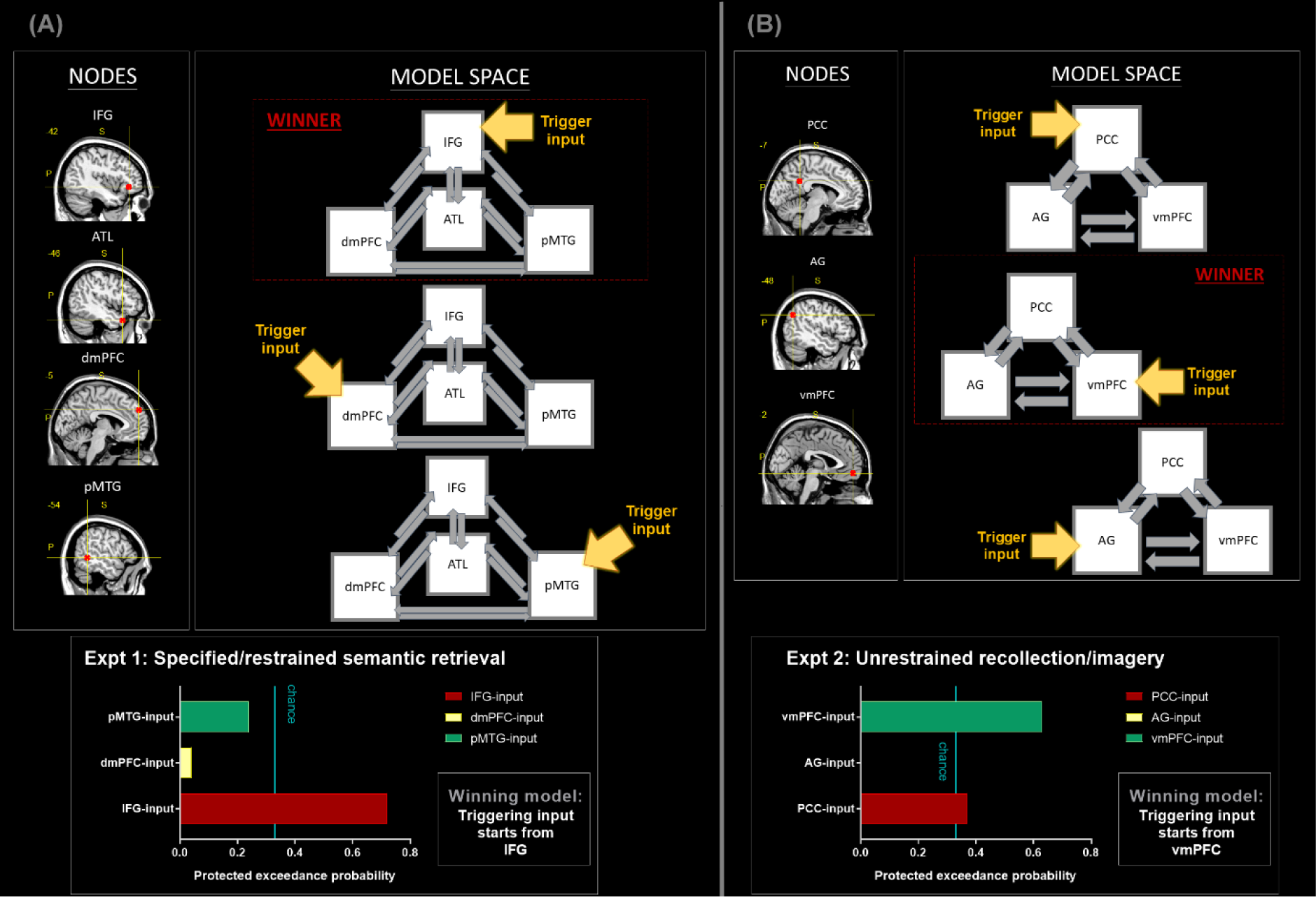
**(A)** DCM-5: All analyses are based on the data of Experiment 1. All of the nodes belong to the SN. **(B)** DCM-6: All analyses are based on the data of Experiment 2. All of the nodes belong to the DN.

## Discussion

In the present study, we conducted a series of neuroimaging analyses to understand the fusion and fission between the DN and SN under various experimental contexts. Our analysis uncovers a highly reliable bipartite split between the two networks. The DN shows a strong antipathy for externally-directed visuospatial tasks, tends to be withdraw from tasks that require attention to external entities (e.g., reading text, pushing buttons), and prefers self-referential processing to other-referential processing even when the psychophysical settings of tasks are yoked. By comparison, the SN exhibits the opposite pattern – it has a moderate dislike for visuospatial tasks, actively participates in various socio-semantic tasks unaffected by the needs to attend external entities, and prefers the other-referential processing to self-referential processing. Furthermore, a task context that encourages participants to focus inwardly drives neural dynamics to arise from the DN (with the vmPFC being the most robust initiating region) whereas a context that requires an outward focus drives neural dynamics to emanate from the SN (with the IFG being the most reliable lead-off area). Taken together, the clearly distinct functional profiles of the DN and SN suggest that, while the two networks similarly show a distaste for tasks requiring non-meaningful sensory-motoric processing, they diverge along an ‘internal *vs.* external’ neurocognitive dimension, with the DN being more inwardly-biased and the SN more outwardly-biased. Below we discuss our findings under the framework of recent proposals concerning a macro-scale gradient that span across the entire cerebral zones.

### Large-Scale Gradational Structure of the Human Cortex

Recent advances in mapping the human cortex have discovered a clear gradient of structural and functional features that spans across the sensory-motoric system and the multimodal system, found in both humans and non-human primates (10, 12, 13, 43-45). An evident neurocognitive dimension of ‘internal *vs.* external preference’ underlies this cortical gradient – on one extreme of the spectrum, the cortical regions are highly modality-specific (e.g., the primary visual/auditory cortex; 10), respond to external events occurring here and now (i.e., showing narrow spatiotemporal receptive fields; 46), and prefers concrete, perceivable stimuli to abstracted, conceptual stimuli (47). Regions on the other extreme of the spectrum (e.g., the core DN areas; 10), on the contrary, exhibit the opposite characteristics – multimodal, able to represent information across long spatial and temporal extents, and sensitive to conceptual-mnemonic stimuli. Juxtaposed between the polar extremes are multiple intermediate zones, such as the SN that leans towards the DN extreme, the dorsal-attention system that leans towards the sensory-motoric extreme (48), and the frontoparietal control system that situated midway between the two ends (49). Within the frontoparietal control system, recent evidence has demonstrated that this middle ground further fractionates into two sectors: one subsystem preferentially coupled with the attention and sensorimotor systems, and the other subsystem preferentially linked with the DN and SN (13; a similar bipartite split was also found in the dorsal attention network, see 47). In the present study, we provide crucial evidence that delineates the fine-grained fractionation within the high-level cerebral territory by revealing the commonalities and, more importantly, the striking distinctions in the functional profiles of the DN and SN. Our findings echo with recent evidence that, while the spatial arrangement of different functional networks is configured in an idiosyncratic manner in each individual’s brain, a clear-cut ‘motif’ (*cf.* Braga *et al.*, 2017) of bipartite split is consistently observed at multiple networks, producing a gradational structure across the entire brain (50).

### Underpinning of the Gradient Structure

Various hypotheses have been proposed regarding the mechanisms that drive the formation of the cortical gradient of the human brain. One potential root cause might be the cytoarchitecture and myeloarchitecture of the cortical sheet (for review, see 43). For instance, it has been shown that cortices of the sensory-motoric system contain higher numbers of neurons but relatively fewer synaptic connections between cells whereas the multimodal system (e.g., the DN or prefrontal cortex) contains fewer neurons but more synapses (51). The level of myelination also differs; sensory-motoric regions are more myelinated (hence, permitting swift conduction of information flow and prompt response to external stimuli) whereas higher-order regions are less myelinated and requires longer time to process (52). In addition, studies based on diffusion tractography (53, 54) and large-scale connectome database (55) have coherently identified a profile of graded connectivity that the SN and DN are transmodal zones where multiple circuitries progressively converge, receiving inputs from various primary-sensory and intermediate zones. Such structural-anatomical evidence suggests a possibility that, although many regions of the SN are often incorporated into the canonical umbrella term ‘default-mode system’, these regions might have different cyto- and myelo-structures and connectivity profiles, which make the SN behaves differently from the core regions of the DN on a variety of tasks as we demonstrate here. One possibility could be that, while both the DN and SN are situated on the multimodal side of the gradient, there is relatively shorter ‘distance’ (in terms of the length of graph metric) from the SN the sensory-motoric cortices. This could make the SN more responsive to external sensory stimuli, compared to the DN that has longer, more convoluted access to sensory cortices. This speculation awaits future investigation.

### Integration between semantic cognition with default-mode processes

Brain regions that support different aspects of semantic cognition have long been studied separately from the regions that constitute the DN, despite the fact that the two systems show substantial overlap (9) and are closely connected (22, 23). Under the framework of a macro-scale cortical gradient, the DN and SN could be construed as two complementary systems that work in tandem to serve high-level cognition. The DN areas have been shown to be maximally distant from the sensory-motoric cortices (10). Its locus explains why the DN sits atop the information processing hierarchy and represents most abstract forms of thoughts divorced from different input modalities (e.g., social relationships or self-related contemplation). In contrast, the location of the SN allows it to interface between the DN, the frontoparietal control system and the sensory-motoric system – which is critical to controlled semantic cognition (14). The formation of generalisable, coherent concepts is known to rely upon the ATL, a transmodal representational hub that can interface with all verbal and sensory-motoric modalities simultaneously (14, 15). In addition, for controlled semantic processing, the semantic representational system needs to interface with executive control mechanisms, underpinned by the IFG and pMTG, in order to generate contextually appropriate behaviours (42). Presumably, the DN and SN work together when the context entails goal-directed cognitive operation that combines self-referential processing with semantic knowledge, as in the retrieval of autobiographical memory.

### Conclusion

In the present study, we report a series of fMRI findings that manifest a clear-cut bipartite split in the functional profiles between the DN and SN, evident both in their response amplitudes to different situations and neural dynamics. These discoveries are consistent with recent theories regarding a macro-scale cortical gradient that encompasses the entire cerebrum. The bipartite split on various functional spectrums (inwardly-directed, self-referential, abstract, multimodal *vs.* outwardly-directed, other-referential, concrete, and unimodal) indicates the crucial need that understanding the relationship between the DN and SN, as well as their relationship with other functional networks, requires more thorough understanding about the macro-scale architecture of the human brain.

## Methods

### Participants

Twenty-four volunteers gave informed consent before their participation. The sample consisted of an 18/6 female-to-male ratio, with average age = 25 years old and SD = 7. All volunteers are right-handed and speak English as their mother tongue. All of them completed the magnetic resonance imaging safety screening questionnaire before the experiment, and reported not having any neurological or psychiatric condition. This study was reviewed and approved by the local research ethics committee.

### Design

Participants completed two experiments in a single session. In Experiment 1, we modified a well-established experimental paradigm that has been widely used to probe the neural basis of self-knowledge (e.g., 7, 29, 56). Participants were asked to complete various tasks under a typical psychophysical context in which they were required to make a trial-by-trial response as quickly as possible within a brief timeframe. There were four conditions: (*i*) The Self-Referential Task: Participants read adjectives describing various personality traits and assessed whether the words suitably describe the characteristics of themselves; (*ii*) The Other-Referential Task: Similar to the Self task, participants read adjectives and assessed whether they suit the Queen Elizabeth’s personality; (*iii*) The Visuospatial Task: Participants viewed a pair of meaningless scrambled visual patterns and answered whether the two were mirror inverse of each other; (*iv*) Mind-Wandering (Rest): Participants passively viewed a blank screen while awaiting the next task block to begin.

Experiment 1 consisted of three runs of scanning. Stimuli were presented using a block design, controlled with *E*-Prime (Psychology Software Tools). Each run was 432-sec in duration, with each of the four conditions (Self/Other/Visuospatial/Rest) having six blocks. The order in which task-conditions were presented was fully counterbalanced across participants so that each task-condition was equally likely to appear in every possible slot of the sequences, with stimuli randomly drawn from a designated stimuli-set (also counterbalanced across participants) for a given scan and shuffled across blocks. Each block contained five trials. Each trial began with a fixation dot (0.8 sec), followed by visual stimuli shown for 2.8 sec and no inter-trial interval. In the Self and Other conditions, we displayed the target of assessment (Self or Queen) above the fixation dot and asked participants to answer whether an adjective word (below the central dot) suitably described the personality of the target individual. In the Visuospatial condition, we displayed two scrambled visual patterns (squiggly lines made from randomly breaking and recombining word text of other conditions); participants performed mental rotation and answered whether the two patterns were left-right flipped. Participants were required to react as fast (and accurately for the Visuospatial Task) as possible within the 2.8-sec limit. In the Rest periods, we displayed a blank screen and instructed participants to stay awake and still while awaiting the next task. Participants reacted to the questions by pressing a button on a MR-compatible response pad with their right hand. All visual stimuli were displayed on a mid-grey background, using a high-resolution LCD goggles (NordicNeuroLab) mounted on top of the head coil.

A total of 180 adjectives were used in the Self and Other conditions, with 90 words used in each condition. The mapping with which a word was shown in either the Self or Other condition was fully counterbalanced across the participants so that each adjective was equally probable to be assessed with reference with self or the Queen. The stimuli-sets contained equal proportion of positive traits and negative traits, evenly allocated to the two conditions. Moreover, we controlled the lexical frequency (based on the British National Corpus) and word length (based on the number of letters; average±SD: 8±2 letters in both conditions) of the stimuli so that the stimuli-sets used in the two task conditions was matched on these psycholinguistic properties.

Experiment 2 consisted of four runs of scanning. We adopted and modified an established experimental design of a landmark study that has been widely used to probe functions of the default network (5). Stimuli were presented using a slow event-related design. Each run was 432-sec in duration. There were four conditions – Autobiographical Memory (AM), Theory of Mind (ToM), Visuospatial Search (VS), and Rest. Each run consisted of 18 active-task events (AM, ToM, VS; six events per condition per run, giving 24 events per condition for the whole experiment) and 18 randomly-jittered Rest intervals intervening between active-task events (duration of jittered Rest – average±SD = 6±2.74 sec, range: 2 – 12.5 sec). The order in which task-conditions were presented was fully counterbalanced across participants so that the events of each task-condition were equally likely to appear in every possible position of the sequences, with stimuli randomly drawn from a designated stimuli-set (also counterbalanced, hence each set being equiprobable to be used in Run 1–4). In the AM task, participants were required to recollect personal experiences (including their own thoughts and feelings at the time, as well as the temporal-spatial contexts) related to the theme of a photograph depicting human activity (see below for details). In the ToM task, participants were required to imagine how the individual(s) in the photograph might be thinking or feeling. In the VS task, participants viewed mosaic scrambled patterns and search for a tiny grey triangle hidden in the patterns.

Each trial began with a cue word for 1 sec, prompting participants the upcoming task (AM: Remember, ToM: Imagine, VS: Search). This was followed by a centrally presented image (600 × 380 pixels), as well as a short text below the image, displayed for 15 sec. In the AM and ToM conditions, the images were photographs depicting people in various situations of life (e.g., seeing a dentist, cooking in the kitchen, protesting in a rally, etc.). The short text below was 20 words in length, serving as a cue to help participants with autobiographical recollection or imagining about others’ thoughts and feelings (e.g., AM: “*Remember the time you learnt the outcome of Brexit referendum. How did you feel? How did you respond to it?*” or ToM: “*Imagine what the girl who’s holding the Christmas cracker is thinking and feeling. Also imagine how her grandpa would respond*”). In the VS condition, the images were made from scrambling images of other conditions into random mosaic patterns, and the text below simply said “*Is there a tiny triangle hidden in the pattern?*” The triangle was present in a half of the trials. After the 15-sec interval during which participants recollected, imagined, or searched, a question was shown for 2 sec asking participants to rate how vivid their memory or imagery was (1: Very vivid, 2: Somewhat vivid, 3: Not at all vivid) by pressing a designated button on the response pad. In the VS condition, the questioned asked if they saw a triangle, and participants answered with a binary ‘yes’ or ‘no’ key response. Participants were instructed to concentrate on the recollection, imagery, and visual search processes during the 15-sec interval, and only make a response when the question was shown at the end.

Prior to scanning, we used a similar 3-step training protocol to that of the Spreng and Grady study (5) to ensure that all of the participants were able to engage confidently in retrieving autobiographical events related to the picture’s topic and imagining the thoughts and feelings that people in the picture might have. Forty-eight photographs depicting human activities and interactions, all of them containing at least one person or more, were used in the AM and ToM conditions, with a half of them used in one condition and the remaining half used in the other. We fully counterbalanced the images and conditions across participants to ensure that: (*i*) each photograph was equally likely to be presented as a cue in the AM and ToM condition, ruling out stimuli-specific effects; (*ii*) for each participant, separate sets of photographs were used as cues for the AM and ToM contexts, thus preventing contamination from the other condition.

### Scanning

Full details of data-acquisition parameters, procedures of pre-processing, and statistical analysis are reported in *SI*. Here we provide the key information: The brain regions of our primary interest are situated in the rostro-ventral aspects of the brain (e.g., the ATL, the vmPFC), which are known to particularly susceptible to signal dropout (57). To combat the dropout issue in these target areas, we adopted a dual-echo EPI sequence, which has been demonstrated to improve signal-to-nose ratio around these rostro-ventral regions relative to other conventional imaging protocols (e.g., 23, 42, 57). A customised procedure was used to combine the two echo-time images of each brain volume. Using SPM8, we integrated the standard pre-processing procedures with multiple correction methods to prevent image distortion and improve inter-participant alignment for group analysis. For statistical analysis at the individual level, all of experimental conditions of each experiment were modelled explicitly as separate regressors, while mind-wandering periods (rest) were modelled implicitly, as per the default of SPM. Blocks/events corresponding to our experimental factors were convolved with a canonical haemodynamic response function. Response-execution periods in Experiment 2 were modelled as a separate regressor. Six motion parameters were entered into the model as covariates of no-interest. Behavioural reaction times were also modelled as parametric modulators to rule out the confounding influences due to task difficulty or cognitive effort.

### Regions of interest (ROIs)

In *SI*, we report the full details of the 10 ROI-coordinates in the MNI stereotaxic space and pinpoint the sites on the template images. All of these ROIs are selected independent of the task contrasts of the two experiments, based on the meta-analysis outcomes of relevant literatures. Specifically, we surveyed four relevant studies of neuroimaging meta-analyses (41, 58-60) about the neural correlates of default-mode functions (the targets include brain area robustly related to *mind-wandering, daydreaming, self-representation, autobiographical memory, theory of mind, episodic memory*), semantic cognition (including *semantic memory, semantic control, conceptual knowledge*), and social cognition (including *theory of mind, mentalising, empathy*). Using the activation likelihood estimates (ALE) identified by these topic-focused meta-analyses, we identified 10 important nodes closely associated with different aspects of default-mode and semantic-related functions – the dmPFC, vmPFC, PCC, bilateral AG, bilateral TPJ, left ATL, left IFG, left pMTG. At each of the ALE peak-coordinates, we created a spherical ROI with 6-mm radius. Next, we evaluated the suitability of the selected ROIs by using the ‘association-test’ function of NeuroSynth (a large-scale fMRI meta-analysis platform; 61). Based on the results of terms-based search (*default-mode, semantic memory, theory of mind, mentalising*) that contain 2,214 studies and 84,573 peaks, NeuroSynth generated the ‘association-test’ maps that represent brain areas statistically significantly (FDR-corrected at *q* < 0.01) associated with these functions. We then compared our selected ROIs and these maps, and confirmed that all of the 10 chosen ROIs are encompassed within the meta-analytic maps (see *SI* for illustrations).

### Dynamic causal modelling (DCM)

We performed a series of six DCM analyses using DCM10 in SPM8 (35). DCM-1, DCM-3, and DCM-5 were performed on the data of Experiment 1, whereas DCM-2, DCM-4, and DCM-6 were done on the data of Experiment 2. For each DCM, we localised the network nodes based on the same coordinates that we defined using relevant literatures, verified using NeuroSynth, and confirmed their involvement in the tasks using a series of ROI analysis. Each network node was localised using a spherical ROI with 6-mm radius centred at the designated coordinate. Activated voxels within each node were identified using the relevant contrasts (i.e., Experiment 1: Self+Other > Rest; Experiment 2: AM+ToM > Rest) thresholded at *p* < 0.001 for voxel intensity. We used the SPM’s default algorithms to extract the first eigenvariate to obtain the time-series of activity, as per the standard procedure of DCM. This process was repeated for all of the selected regions in each participant. All of the DCM were set to have one state per region and without stochastic modulatory effect. Based on the extracted activity, we constructed and estimated DCM separately for each participant, and then employed the Variational Bayesian Analysis (VBA) Toolbox (39, 40) to conduct random-effects (RFX) group analyses using its functions of Bayesian models selection. The RFX procedure outputted the protected exceedance probability (PEP; 39), which is a unbiased probabilistic estimate that quantifies how likely that, for an individual randomly drawn from the population, the model being considered provides the best fit to this person’s fMRI data than any other model, above and beyond the chance-level. The chance-level represents the probability of all models being equally likely (1/K, with K being the number of models; we had 0.33 as the chance-level for all analyses). Bayesian model selection does not rely on a binary, arbitrary cut-off threshold. Rather, it relies on identifying the model that shows the highest, sufficiently above-chance PEP.

Here we specify the model spaces: (*i*) In DCM-1 and DCM-2, we focused on the relationship between the dmPFC and the vmPFC; for each DCM, we constructed three models, and each model contained two nodes, with the input signal entering via the dmPFC, the vmPFC, or both nodes. The two nodes were set to be mutually connected. (*ii*) In DCM-3 and DCM-4, we focused on the relationship amongst the IFG, the ATL, and the vmPFC; for each DCM, we constructed three models, and each model contained three nodes, with the input signal entering via the IFG, the ATL, and the vmPFC. All of the three nodes were set to be mutually connected. (*iii*) In DCM-5, we focused on the relationship between nodes within the SN (the IFG, the pMTG, the dmPFC, and the ATL) and tested only using Experiment 1’s data. For each DCM, we constructed three models, and each model contained four nodes, with the input signal entering via the IFG, the pMTG, and the dmPFC. We did not include the ATL as an entry node here because the outcome of the preceding DCM-3 had already indicated that the probability of ATL being the winner was close to zero. In order to constrain the size of model space as three models and maintain a consistent chance-level across analyses, we included only the IFG-, pMTG, and dmPFC-input models. All of the four nodes were set to be mutually connected. (*iv*) In DCM-6, we focused on the relationship between nodes within the DN (the PCC, the AG, the vmPFC) and tested only using Experiment 2’s data; for each DCM, we constructed three models, and each model contained three nodes, with the input signal entering via the PCC, the AG, and the vmPFC. All of the three nodes were set to be mutually connected.

## Acknowledgements

This research was funded by an MRC programme grant to MALR (MR/R023883/1) and a Sir Henry Wellcome Fellowship (201381/Z/16/Z) to RC.

