## Supplemental information for "Bipartite functional fractionation within the default network supports disparate forms of internally oriented cognition"

Rocco Chiou ✉, Gina F. Humphreys, & Matthew A. Lambon Ralph ✉
MRC Cognition and Brain Sciences Unit, University of Cambridge, UK

✉ Correspondence to this work can be addressed to either of the authors by email:
(**RC:**, **MALR:**)

### Supplemental Methods

MRI acquisition. All scans were acquired using a 3T Phillips Achieva scanner equipped with a 32-channel head coil and a SENSE factor of 2.5. A dual-echo EPI sequence was used to maximise signal-to-noise ratio in the rostral-ventral surface of the brain that has been known to susceptible to signal dropout (1). Using this technique, each scan consisted of two images acquired simultaneously with different echo times: a short echo optimised to obtain maximum signal from the ventral parts and a long echo optimised for whole-brain coverage. The sequence included 31 slices covering the whole brain with repetition time (TR) = 2.8 sec, short / long echo times (TE) = 12 / 35 ms, flip angle = $85^o$, field of view (FOV) = 240 × 240 mm, resolution matrix = 80 × 80, slice thickness = 4 mm, and voxel dimension = 3 × 3 mm on the *x*- and *y*-axis. To reduce ghosting artefacts in the temporal lobes, all functional scans were acquired using a tilted angle, upward $45^o$ off the AC-PC line. Functional scans of the two main experiments were collected over seven runs; each run was 432-sec long during which 155 dynamic scans were acquired (alongside 2 dummy scans, discarded). To tackle field-inhomogeneity, a $B_0$ field-map was acquired using identical parameters to the functional scans except for the following: TR = 599 ms, short / long TEs = 5.19 / 6.65ms. Total B0 scan time was 1.6 minutes. A high-resolution $T_1$-weighted structural scan was acquired for spatial normalisation, including 260 slices covering the whole brain with TR = 8.4 ms, TE = 3.9 ms, flip angle = $8^o$, FOV = 240 × 191 mm, resolution matrix = 256 × 163, and voxel size = 0.9 × 1.7 × 0.9 mm. Total structural scan time took 8.19 minutes.

Pre-processing. Analysis was carried out using SPM8 (Wellcome Department of Imaging Neuroscience). The functional images from the short and long echoes were integrated using a customised procedure of linear summation (1, 2). The combined images were realigned using rigid body transformation (correction for motion-induced artefacts) and un-warped using B0 field-map (correction for field-inhomogeneity). The averaged functional images were then co-registered to each participant's $T_1$ anatomical scan. Spatial normalisation into the MNI standardised space was achieved using the DARTEL Toolbox of SPM (3), which has been shown to produce highly accurate inter-subject alignment (4). Specifically, the $T_1$-weighted image of each subject was partitioned into grey-matter, white-matter, and CSF tissues using SPM8's 'Segmentation' function; afterwards, the DARTEL toolbox was used to create a group template derived from all participants. The grey-matter component of this template was registered into the SPM grey-matter probability map (in the standard MNI stereotactic space) using affine transformation. In the process of creating the group's template brain using individual $T_1$, for each individual DARTEL estimated 'flow fields' that contained the parameters for contorting native $T_1$-weighted images to the group template. SPM8 deformation utility was then applied to combine group-to-MNI affine parameters with each participant's 'flow fields' to enable tailored warping into the MNI space with better accuracy. The functional images were then resampled to a 3 × 3 × 3 mm voxel size. Smoothing was subsequently applied using an 8-mm Gaussian FWHM kernel, consistent with prior studies (e.g., 1, 5).

*GLM analysis*. For each participant, contrasts of interest were estimated using general linear model (GLM) convolving the experimental design matrices with a canonical haemodynamic response function, with resting periods modelled implicitly. Motion parameters were entered into the model as covariates of non-interest. For Experiment 2, we separately modelled the events of interest (i.e., the 15-sec interval of AM, ToM, and VS) and the button-response interval at the end so that the results are not contaminated by response preparation or execution. Moreover, we included each participant's reaction time (RT) of all active-task performance as parametric modulators, allowing us to rule out any brain activation driven by task difficulty or cognitive effort when assessing the effects of experimental manipulation. Low-frequency drifts were removed using a high-pass filter of 128 sec. Contrast images from the individual-level analyses were then submitted to random-effect models in the group-level analyses.

*Coordinates of the regions of interest (ROIs)*. The table below displays all of the ROIs used in the present study, their coordinates in the MNI stereotaxic space, and location pinpointed on the MNI template by the yellow crossbar. Also rendered on the template is the meta-analysis outcomes, based on the NeuroSynth database, of brain regions related to default-mode network and semantic memory.

| ROI | MNI-coordinate | Location |
|---|---|---|
| dmPFC | -5, 54, 34 | 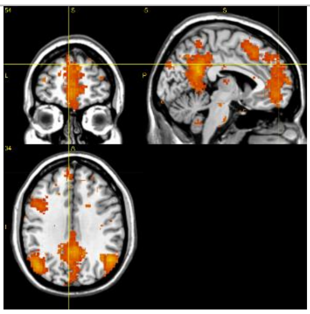 |
| vmPFC | -2, 54, -12 | 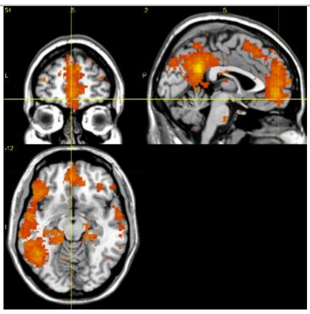 |
| PCC | -7, -48, 31 | 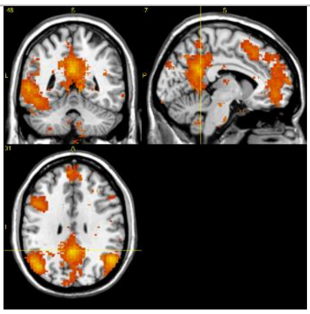 |
| L.AG | -48, -64, 34 | 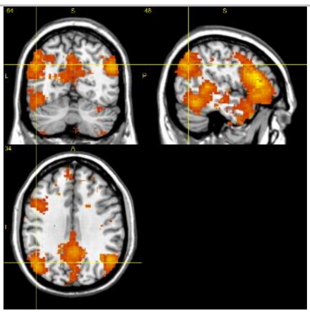 |
| R.AG | 48, -64, 34 | 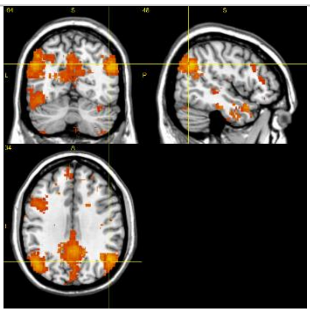 |

| | | |
|---|---|---|
| ATL | -46, 18, -26 | 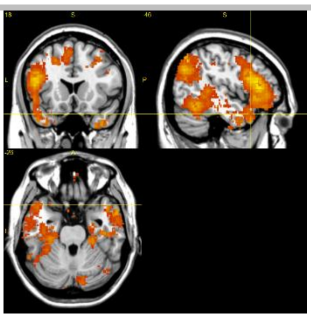 |
| IFG | -42, 34, -6 | 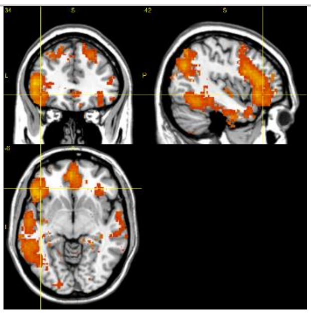 |
| pMTG | -54, -49, -1 | 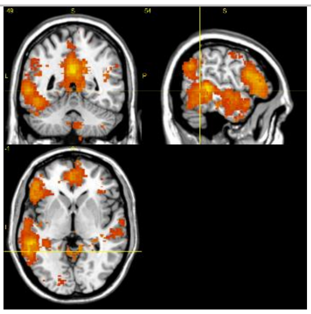 |
| L.TPJ | -57, -42, 20 | 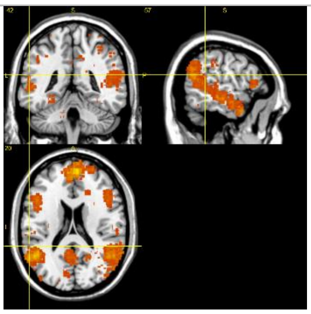 |
| R.TPJ | 57, -42, 20 | 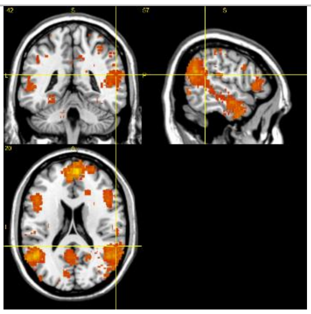 |

**Supplemental Results**

*Behavioural analysis*. In Experiment 1, participants showed comparable reaction times for the Self- (average±SD: 1071±35 ms) and Other-Referential (1056±39 ms) tasks, with no statistically reliable difference between them (paired *t*-test, $p = 0.42$). In Experiment 2, participants also showed comparable reaction times for the AM (807±36 ms) and ToM (832±44 ms) conditions, with them having no reliable difference ($p = 0.39$; note that these reaction times represent the latency to rate vividness *after*, rather than during, an AM- or ToM-interval). More importantly, the results of vividness ratings indicated that participants were able to engage in highly vivid recall and imagery (AM: 1.25±0.04; ToM: 1.28±0.05, both approached the ceiling-level on the given scale), while the ratings did not differ between contexts ($p = 0.64$). Together, these data showed that, in both experiments, the tasks specifically designed to probe the functionality of the default network are matched on the general cognitive effort required (as indicated by reaction times) and the clarity of introspective experiences (as indicated by the vividness ratings). Also note that in all fMRI analysis, reaction times (including those of the visuospatial tasks) were included as additional regressors to factor out their influences on the neural data.

Experiment 1: Self-representation in the ventromedial prefrontal cortex (vmPFC). It has been repeatedly demonstrated that the vmPFC amplifies its activity when participants evaluate personality descriptions with reference to self (e.g., 6, 7, 8). However, after a closer inspection at these results, we found that such an effect was actually driven by *less deactivation* for the '*Self > Rest*' contrast (which often results in no difference), compared to other contrasts that induce *significant deactivation* (e.g., '*Other > Rest*' or '*Letter-case > Rest*', due to greater mPFC activity during rest). In this supplemental analysis of Experiment 1, we further investigated this issue by comparing the Self condition with three different baselines (Other, Visuospatial, and Rest). All of the analyses were conducted using whole-brain interrogation, thresholded at $P < 0.05$ FWE-corrected for voxel intensity. As clearly illustrated in Fig. *S*1, the vmPFC responds the self-evaluation task and rest (mind-wandering) with comparable activation level, resulting in no significant cluster in the vmPFC region in both contrasts ('*Self > Rest*' & '*Rest > Self*'; Fig. *S*1 right). This is consistent with previous finding that participants tended to think about themselves during the resting period (9), driving vmPFC activity to persist despite no task during rest. However, a significant vmPFC-cluster emerges when we searched for '*Self > Other*' and '*Rest > Other*' (Fig. *S*1 middle). The size of this cluster further expands when we searched for '*Self > Visuospatial*' and '*Rest > Visuospatial*' (Fig. *S*1 left). This suggests a gradational pattern – the vmPFC is least involved during the visuospatial task, most involved during the Self task and mind-wandering, situated in between during the Other task.

Taken together, these data corroborate and complement our findings reported in the main article – when the contrast baseline is rest (which is known to full of self-referential thoughts), a more inwardly-oriented autobiographical memory task enhances vmPFC activity beyond its level during resting-state, whereas a more outwardly-oriented self-traits task tended to induced vmPFC activity commensurate with its resting-state level. This echoes our argument made in the main text that while the involvement of the vmPFC in various cognitive processes is sensitively modulated both by the 'inward *vs*. outward' factor both at a more physical level (attending external stimuli *vs*. attending internal thoughts) and an abstract level (thoughts about self *vs*. thoughts about other).

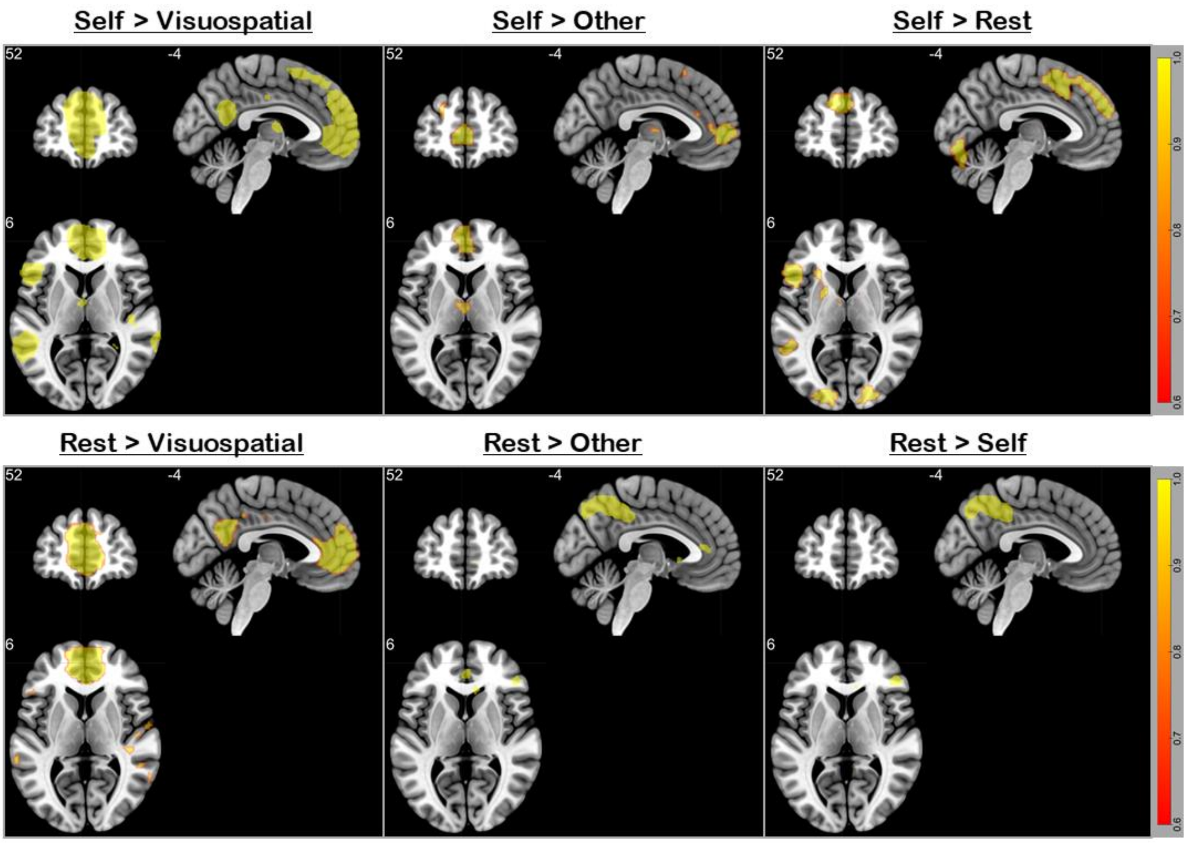

***Figure S1***. All contrasts were thresholded at $p< 0.05$ (FWE-corrected for whole-brain voxels). The upper row illustrates the contrast of (left to right) 'self-knowledge > visuospatial task', 'self-knowledge > other-referential knowledge', and 'self-knowledge > mind-wandering (rest)'. Note that the choice of baseline affects the size of vmPFC cluster, with the rest-baseline engaging the vmPFC most, the visuospatial-baseline engaging the vmPFC least, and the other-baseline being intermediate. A highly similar pattern is found in the contrast of (lower-row, left to right) 'Rest > Visuospatial', 'Rest > Other', and 'Rest > Self', indicating that mind-wandering (rest) equally engages the vmPFC as self-knowledge.

*Experiment 1 & 2: Social-knowledge representation in the dorsomedial prefrontal cortex (dmPFC)*. Unlike the vmPFC that is preferentially engaged when participants process information with reference to themselves, the dmPFC has been found to be ubiquitously active in a wide variety of social contexts. This is also observed in our experiments: As shown in Fig. *S*2, the dmPFC is active both when participants evaluate the personality traits about themselves or the Queen (Experiment 1), and when they retrieve the memories about self-life events and make inferences about the mental states of other people (Experiment 2). This is consistent with previous proposal that the dmPFC plays a more general role in any social situation that entails evaluation of traits, feelings, beliefs, of a social being (e.g., 10).

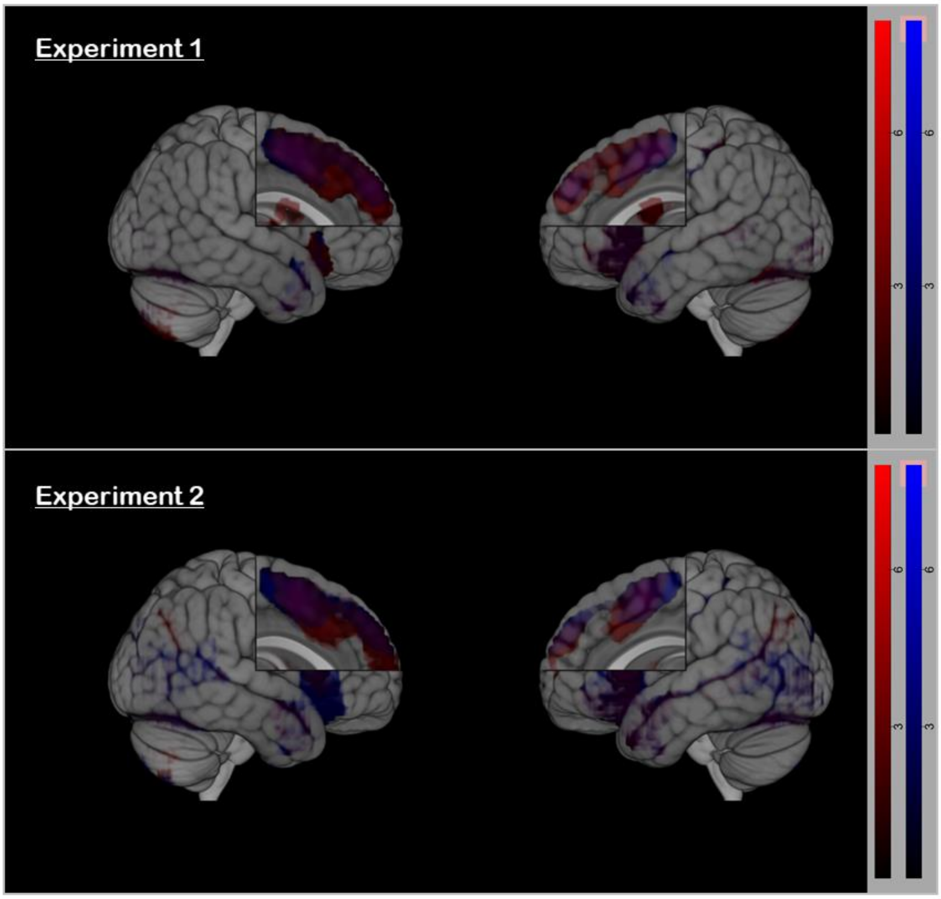

***Figure S2***. All contrasts were thresholded at $p< 0.05$ (FWE-corrected for cluster size) and $p < 0.001$ (for cluster-defining voxel intensity), compared against the resting baseline. Upper row (Experiment 1): Red colour stands for knowledge about self-traits; blue colour stands for other's traits; purple is the conjunction. Lower row (Experiment 2): Red colour stands for autobiographical memory; blue colour stands for theory-of-mind; purple is the conjunction.

Experiment 1 & 2: Frontoparietal activity driven by the visuospatial tasks. We observed highly similar patterns of neural activity driven by the mental rotation task (Experiment 1) and the visual search task (Experiment 2). These two visuospatial tasks activated a set of widely distributed 'task-positive' frontoparietal regions known to underpin spatial attention and executive control. All of the analyses here were examined using whole-brain interrogation, contrasted against the rest-baseline, thresholded at $p < 0.05$ FWE-corrected for voxel intensity. As clearly illustrated in Fig. *S*3, both tasks elicited robust activity of the frontal and parietal regions known to be involved in attention, executive control, and visuospatial working memory (for review, see 11, 12), as well as the bilateral visual cortices.

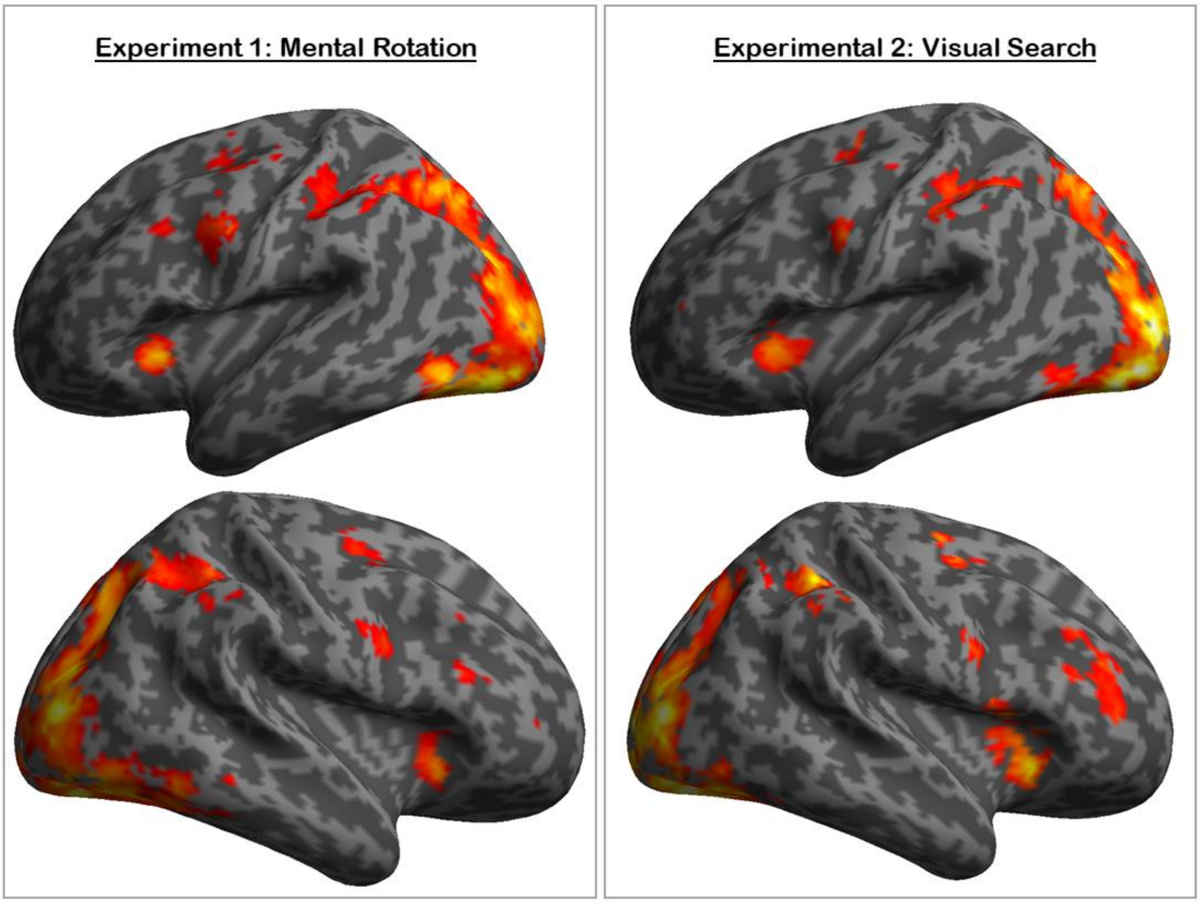

***Figure S3***. All contrasts were thresholded at $p< 0.05$ (FWE-corrected for whole-brain voxels). Left (Experiment 1): The mention rotation task > Rest. Right (Experiment 2): The visual search task > Rest. A consistent pattern of frontoparietal activity, typically seen for demanding visuospatial tasks, is observed in both experiments.
